# The [PSI+] prion and HSP104 modulate cytochrome *c* oxidase deficiency caused by deletion of COX12

**DOI:** 10.1101/2021.10.08.463630

**Authors:** Pawan Kumar Saini, Hannah Dawitz, Andreas Aufschnaiter, Jinsu Thomas, Amélie Amblard, James Stewart, Nicolas Thierry-Mieg, Martin Ott, Fabien Pierrel

**Affiliations:** Univ. Grenoble Alpes, CNRS, UMR 5525, VetAgro Sup, Grenoble INP, TIMC, 38000 Grenoble, France; Department of Biochemistry and Biophysics, Stockholm University, Stockholm 10691, Sweden; Max Planck Institute for Biology of Ageing, Joseph-Stelzmann-Str. 9b, 50931 Cologne, Germany; Wellcome Centre for Mitochondrial Research, Biosciences Institute, Faculty of Medical Sciences, Newcastle University, Newcastle Upon Tyne, NE2 4HH, United Kingdom; Department of Medical Biochemistry and Cell Biology, University of Gothenburg, Gothenburg 40530, Sweden

**Keywords:** Cytochrome *c* oxidase, Cox12, respiration, mitochondria, experimental evolution, Hsp104, prion, PSI+, yeast

## Abstract

Cytochrome *c* oxidase is a pivotal enzyme of the mitochondrial respiratory chain, which sustains bioenergetics of eukaryotic cells. Cox12, a peripheral subunit of cytochrome *c* oxidase, is required for full activity of the enzyme, but its exact function is unknown. Here, experimental evolution of a *Saccharomyces cerevisiae* Δ*cox12* strain for ~300 generations allowed to restore the activity of cytochrome *c* oxidase. In one population, the enhanced bioenergetics was caused by a A375V mutation in the AAA+ disaggregase Hsp104. Deletion or overexpression of Hsp104 also increased respiration of the Δ*cox12* ancestor strain. This beneficial effect of Hsp104 was related to the loss of the [*PSI^+^*] prion, which forms cytosolic amyloid aggregates of the Sup35 protein. Overall, our data demonstrate that cytosolic aggregation of a prion impairs the mitochondrial metabolism of cells defective for Cox12. These findings identify a new functional connection between cytosolic proteostasis and biogenesis of the mitochondrial respiratory chain.

## INTRODUCTION

Mitochondrial energy conversion is pivotal for cellular metabolism and bioenergetics. Energy conversion relies on a series of multi-subunit enzymes, which together form the respiratory chain. Electrons are extracted from reduced metabolic intermediates, originating mostly from the TCA cycle, and transported with the help of mobile electron carriers to the terminal enzyme, cytochrome c oxidase (CcO), which reacts these electrons with molecular oxygen. The chemical energy from the reduced metabolites is thereby converted into proton motive force, which drives ATP formation in mitochondria. In the yeast *Saccharomyces cerevisiae*, the metabolism of so-called respiratory carbon sources (like glycerol, acetate, lactate, ethanol) relies on mitochondrial respiration, whereas glucose can be processed independently via glycolysis.

The biogenesis of the respiratory chain necessitates expression of nuclear and mitochondrial genes. Nuclear-encoded proteins are synthesized in the cytoplasm and are both, post- and co-translationally imported into the organelle by the help of sophisticated import complexes ^1^. Inside mitochondria, the different complexes are formed by the step-wise addition of subunits, which in turn need to be equipped with redox co-factors aiding in electron transfer. This assembly process and the fact that nuclear gene expression needs to be synchronized with mitochondrial translation ^2,3^, makes biogenesis of respiratory chain complexes intricate.

The mitochondrial CcO is the terminal enzyme of the respiratory chain and contains 14 protein subunits, of which three, Cox1, Cox2 and Cox3, are typically encoded in the mitochondrial DNA. These three subunits are conserved from the alpha proteobacterial ancestor of mitochondria. Interestingly, only two of these subunits, Cox1 and Cox2, contain redox co-factors (*a* and *a3* hemes, Cu_A_ and Cu_B_) necessary for electron transfer ^4^, while Cox3 supports the enzyme by establishing a pathway for oxygen diffusion to the active site. Why the mitochondrial enzyme contains 11 additional subunits is currently not fully understood. A few of the subunits have been implicated in regulation of CcO, while others might confer stability to the catalytic core that becomes destabilized because of the comparably rapid rates by which mutations occur in mitochondrial genomes ^5^. Work in *S. cerevisiae* has shown that absence of distinct subunits leads to different consequences, ranging from complete absence of respiration to very small changes in enzyme activity.

In particular, the deletion of Cox12, a peripheral subunit of CcO that resides in the intermembrane space ^6^, drastically decreased the activity of CcO despite hemes *a* + *a3* being detected at ~50% of wild-type (WT) levels ^7^. Interestingly, Cox12 is probably not essential for CcO activity *per se* since detergent-purified CcO remained active despite the loss of Cox12 during the purification ^7^. Yeast Cox12p is the ortholog of human COX6B and two missense mutations in the COX6B1 isoform caused severe clinical symptoms ^8^, supporting the functional importance of the protein. The precise function of Cox12 is currently unknown but Ghosh *et al.* suggested a link with the copper delivery pathway to Cox2 ^9^.

Here we used experimental evolution to alleviate the CcO deficiency of Δ*cox12* cells and we identified the Hsp104 disaggregase as a modulator of the growth defect on respiratory carbon sources. We further demonstrated an unexpected connection between the cytosolic [*PSI^+^*] prion and the functionality of CcO in Δ*cox12* cells. Together, our data reveal a new link in the intricate network that connects the cytosol and mitochondria.

## RESULTS

### Experimental evolution yields Δ*cox12* cells able to grow on respiratory medium

Δ*cox12* cells are deficient for CcO activity, and as such they are unable to grow on respiratory media but can propagate on fermentable medium containing glucose ^7^. We used experimental evolution to select mutations that would rescue the respiratory growth defect of Δ*cox12* cells. To this end, we propagated Δ*cox12* cells in a medium that contained a limiting amount of maltose (as a fermentable carbon source) and an excess of lactate, glycerol and acetate (LGA, as respiratory carbon sources). In those cells, we deleted the *MSH2* gene in order to increase the nuclear mutation rate ^10,11^, and thus accelerate the emergence of evolved phenotypes. The cells were cultured in two replicates (A and B) and were serially transferred 43 times, representing approximately 300 generations (Fig. 1A). After transfer number 8, maltose was omitted from the growth medium as the cultures were able to metabolize LGA. The density of the cultures increased over passages (Table S1), showing that cells improved their respiratory capacities. At the end of the evolution experiment, populations A and B showed robust growth on LGA medium, unlike the Δ*cox12* ancestor (Fig. 1B). We isolated three clones from each population and assayed their growth on media with various carbon sources (Fig. 1C). All evolved clones grew robustly on LGA medium and also on media that contained glycerol-acetate, ethanol or lactate as sole carbon sources (Fig. 1C), supporting that their mitochondria harbored a functional CcO. As the phenotypes of the three tested clones were comparable (Fig. 1C), we selected a single clone per population for further study (Ac and Ba). Interestingly, clone Ba was able to grow on glycerol medium at 37°C unlike clone Ac (Fig. 1D), suggesting that rescue mechanisms might differ in evolved populations A and B.

**Figure 1:**
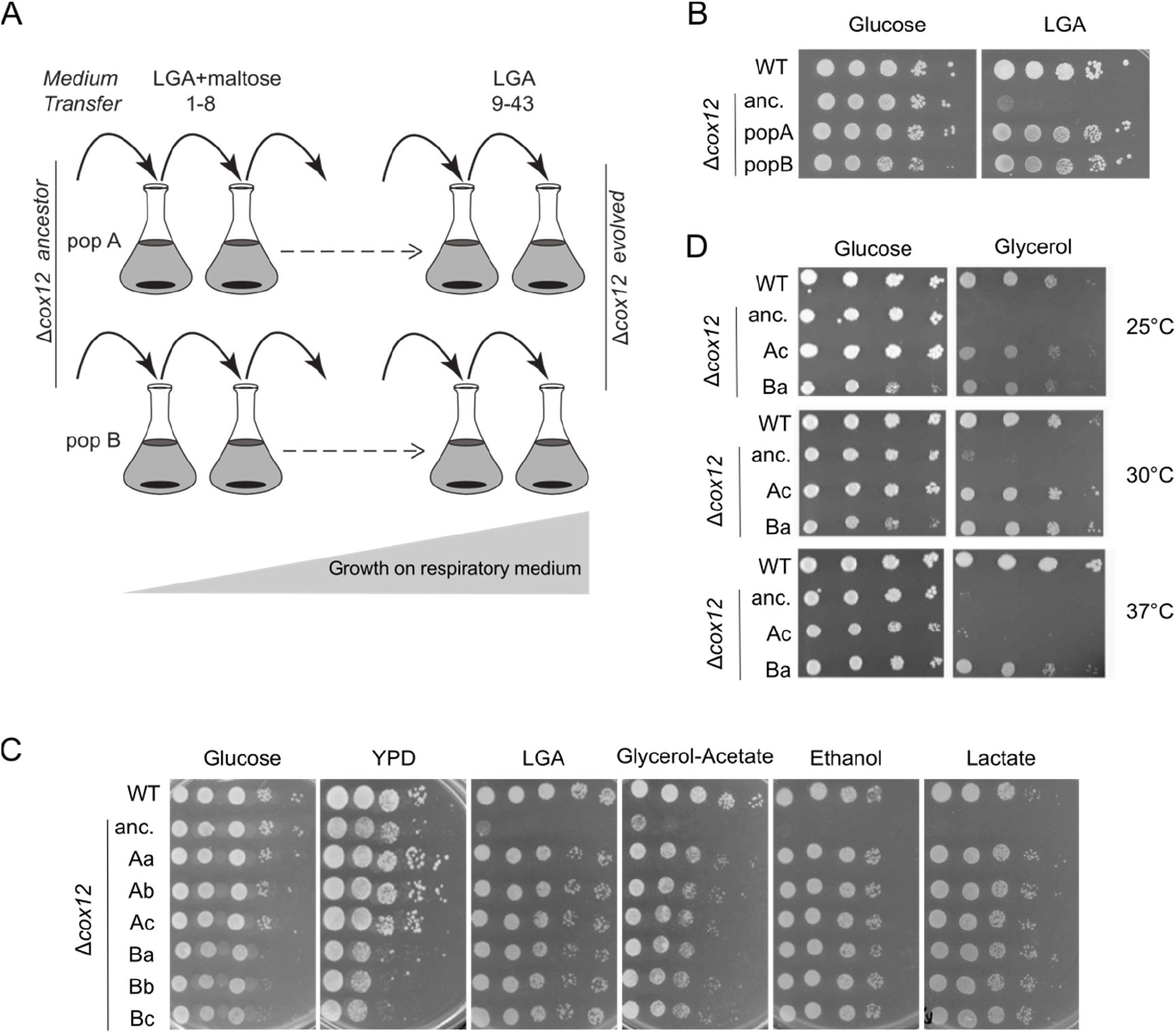
Phenotypic analysis of evolved Δ*cox12* cells. A) Schematic representation of the evolution experiment. B) 10-fold serial dilutions of control strains (WT and Δ*cox12* ancestor) and of populations A and B after transfer 43. The synthetic media contained either glucose, or lactate-glycerol-acetate (LGA) as in the evolution experiment. C) Serial dilution of control strains (WT and Δ*cox12* ancestor) and of clones a-c isolated from populations A and B. The plates contained either YPD medium or synthetic medium supplemented with the indicated carbon sources. The plates were imaged after 2 days (YPD and Glucose) or 4 days at 30°C. D) The indicated strains were tested for growth at various temperatures on synthetic medium containing glucose or glycerol. The plates are representative of 2 independent experiments (B-D).

Since the evolution experiment was conducted with a *msh2::CaURA3* hypermutator strain, the cells accumulated many mutations over 300 generations. Among these mutations, only a few, called here causal mutations, are likely to mediate recovery of respiratory growth in the evolved strains. To characterize the genetic properties of the causal mutation(s), we crossed clones Ac and Ba with the Δ*cox12-HM* strain of opposite mating-type. The Ba diploid did not grow on LGA medium whereas the Ac diploid did (Fig. S1A), indicating that clones Ba and Ac carried recessive and dominant causal mutations, respectively. The Ba diploid was unable to sporulate, consistent with the requirement of a functional mitochondrial respiratory chain for sporulation ^12^. Thus, to evaluate the number of causal mutation(s) in Ac and Ba clones, we crossed them to strain CEN-HM carrying the wild-type *COX12* gene and we analyzed the meiotic progeny by tetrad dissection. As expected, the Δ*cox12* deletion segregated 2:2 and 50% of the Δ*cox12* spores were able to grow on LGA (Table S2), supporting the existence of a single causal mutation in the parental clones Ac and Ba. We analyzed the phenotype of two LGA-positive meiotic clones (P1, P2) and found that it was comparable to that of their respective parents, Ac and Ba (Fig. S1B). Clones Ac-P1 and Ba-P2 were selected for further study as they had inherited the wild-type *MSH2* gene after backcross and thus had a normal mutation rate.

### Δ*cox12* evolved clones compensate reduction of CcO protein levels and activity

Next, we assessed if the restoration of respiratory growth in the evolved clones was related to altered proteins levels of CcO subunits. For that purpose, we investigated yeast cells grown either in fermentable glucose or in galactose, as a respiratory carbon source. As expected, we observed a severe reduction of all analyzed CcO proteins in the Δ*cox12* deletion strain. Interestingly, this effect was largely reverted in clones Ac-P1 and Ba-P2 (Fig. 2A), which showed increased levels of Cox1, Cox2 and Cox13. To evaluate CcO activity, we recorded heme spectra (Fig. 2 B-D) and normalized oxygen consumption to the amount of CcO molecules. The massive reduction of CcO activity in the Δ*cox12* strain (approx. 25% of WT) was alleviated in the evolved Ac-P1 and Ba-P2 clones (approx. 60% of WT, Fig. 2E). From these results, we concluded that the evolved clones Ac-P1 and Ba-P2 recovered respiratory growth because of a causal mutation that increases CcO activity despite the absence of Cox12.

**Figure 2:**
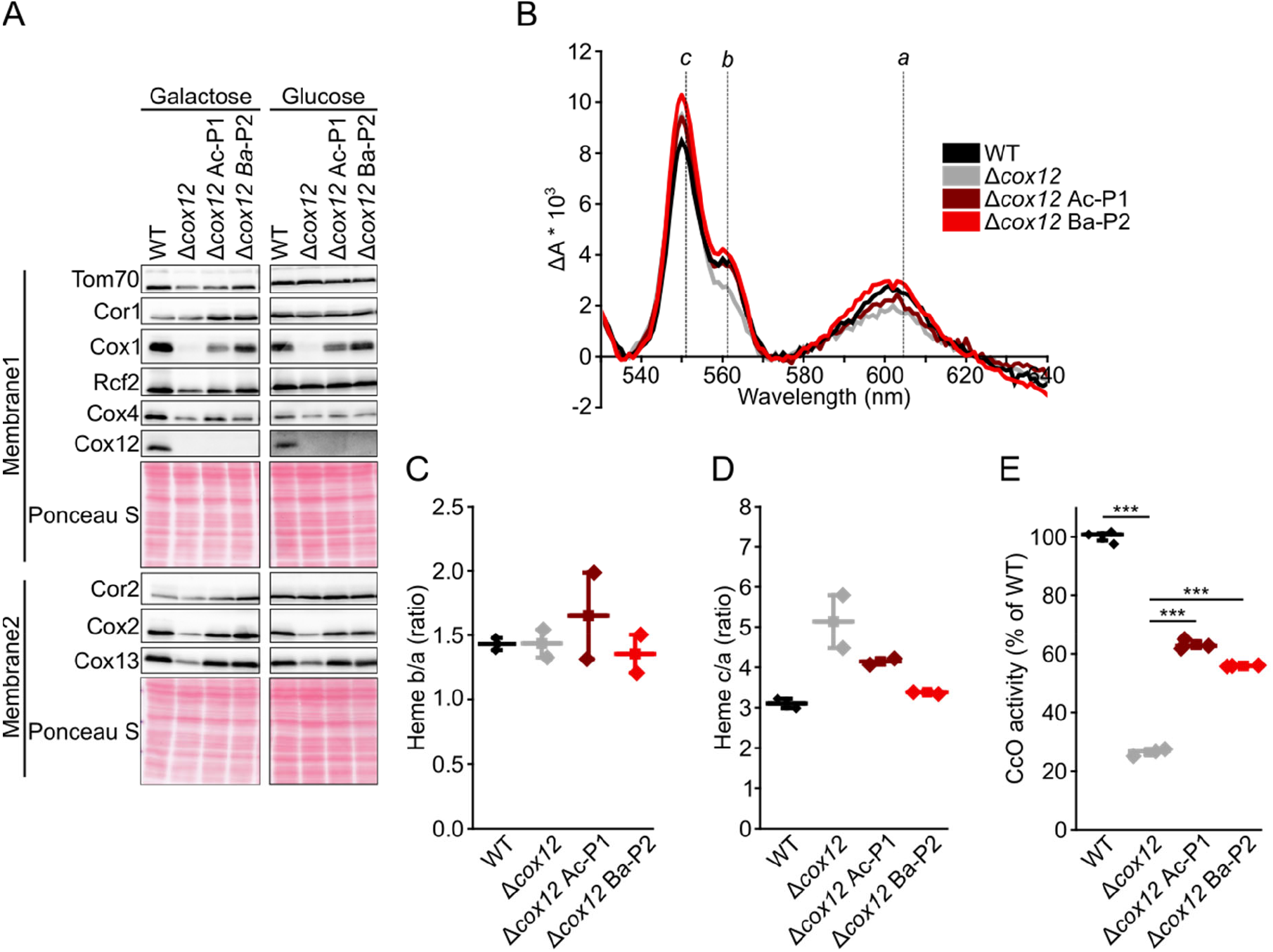
CcO protein levels and activity. A) Steady state levels of indicated proteins were analyzed in depicted strains during exponential growth in YP medium containing either galactose or glucose as carbon source. Ponceau S staining of respective membranes was performed as loading control. B-D) Heme spectra and heme ratios recorded from mitochondria isolated from indicated strains (n=2). E) CcO activity was measured as oxygen consumption in mitochondrial lysates as described in material and method section (n=3). 10 mmol CcO were used for each measurement, as quantified via heme content (B-D). Data was subsequently normalized to CcO activity from wild type mitochondrial extracts. Analyses in C-E are depicted as mean (square) ± s.e.m., median (center line) and single data points (diamonds). Data in E was statistically analyzed by a One-Way ANOVA followed by a Tukey Post Hoc test. Single main effects are depicted as ***P < 0.001

### HSP104 A375V is the causal mutation in evolved Δ*cox12* clones isolated from population B

In order to identify the causal mutations in clones Δ*cox12* Ac and Δ*cox12* Ba, we used an approach that consists in sequencing the genome of clones selected phenotypically within a meiotic progeny, as originally described by Murray and colleagues ^13^. In our case, the causal mutations of interest would confer growth on LGA to evolved Δ*cox12* clones. Thus, we reasoned that in the meiotic progeny of evolved Δ*cox12* clones Ba and Ac crossed to the CEN-HM strain, all Δ*cox12* clones able to grow on LGA (P clones = Positive for growth) should contain the causal mutation, whereas those unable to grow on LGA (N clones = Negative for growth) should contain the wild-type allele. In contrast, non-causal mutations should distribute evenly in pools of P clones or N clones that we constituted from 10 to 15 individual Δ*cox12* clones selected in the meiotic progeny. We sequenced these pools at ~100 X coverage, together with the ancestor Δ*cox12* strain, the wild-type (WT) strain, and individual clones from the meiotic progeny able to grow on LGA (Ac-P1, Ba-P2, see Fig. S1B). In order to identify the causal mutation, we filtered the sequencing results for mutations present in the Ba-P1 clone and Ba-P pool, but absent in the WT, in the Δ*cox12* ancestor and in the Ba-N pool. This analysis yielded three candidate mutations (Table S3). Two of them were positioned in intergenic regions, the third (XII:89746 C→T substitution) was present in the coding sequence of the *HSP104* gene, and this *hsp104*-C1124T caused a A375V missense variant in the Hsp104 protein. A similar analysis did not yield any candidate causal mutation for clone Ac, which was therefore not further considered in this study.

Interestingly, Sanger sequencing revealed that the *hsp104*-C1124T mutation was also present in Δ*cox12* Bb and Bc (Fig. S2A), two other clones originally isolated from population B (Fig. 1C). Hsp104 is a heat shock protein that is induced under various stresses. It exhibits disaggregase activity in cooperation with Ydj1p (Hsp40) and Ssa1p (Hsp70) to refold and reactivate denatured, aggregated proteins ^14^. *S. cerevisiae* Hsp104 counts 908 residues and the A375V mutation lies in the nucleotide binding domain 1 (Fig. S2B), which provides most of the ATPase activity necessary to drive protein disaggregation ^15^. This region shows a high level of conservation and A375 is strictly conserved in sequences from distant species (Fig. S2C). To test if the A375V mutation affected the function of Hsp104, we evaluated its capacity to promote induced thermotolerance, as described previously ^16,17^. As seen in figure 3A, the Δ*hsp104* strain showed a decreased survival at 50°C compared to the WT strain, in agreement with previous reports ^16,17^. Chromosomal integration of the *hsp104*-C1124T gene encoding Hsp104-A375V restored thermotolerance similar to WT (Fig. 3A), showing that the A375V allele did not alter the disaggregase activity.

**Figure 3:**
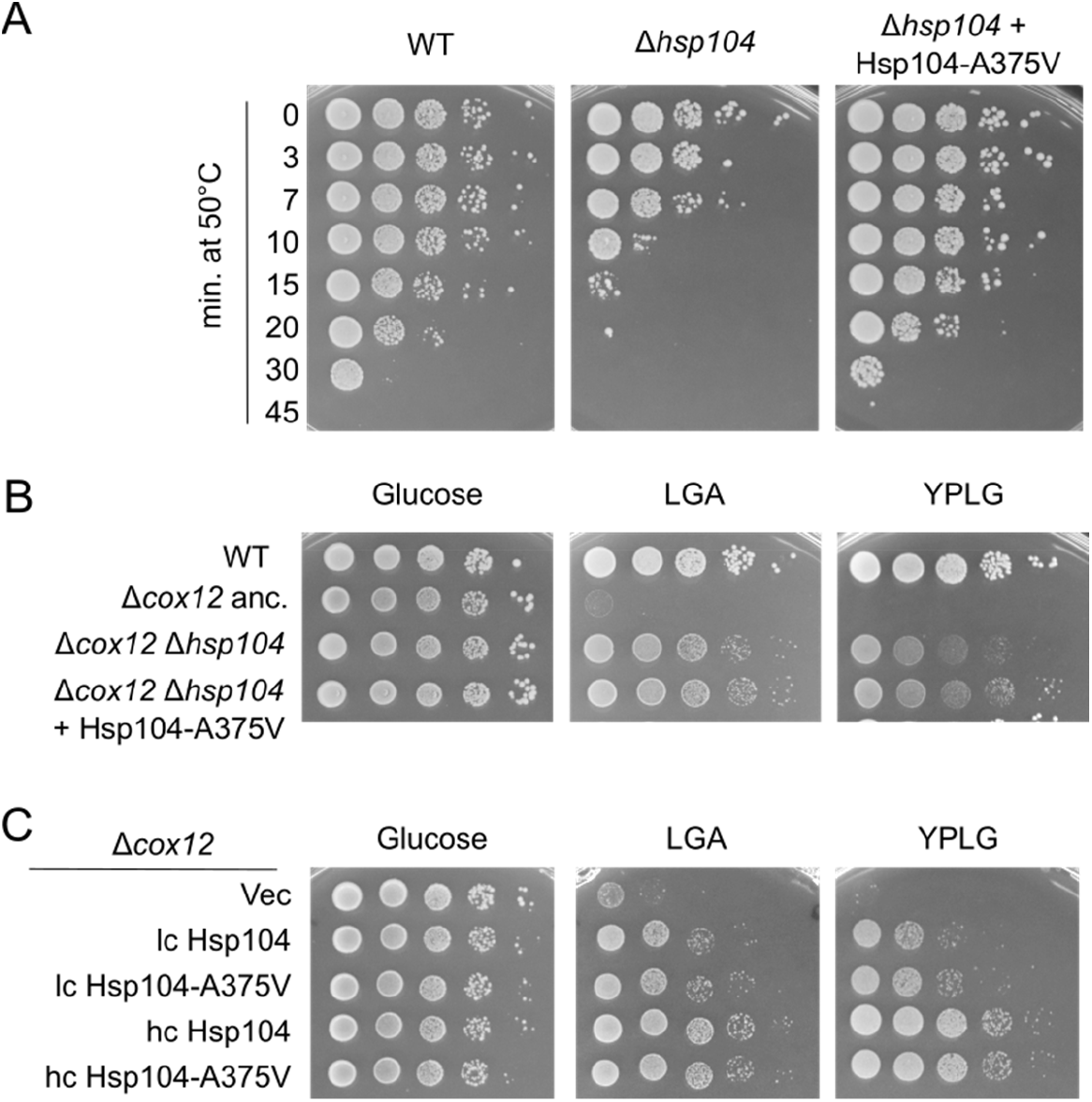
Deletion or overexpression of HSP104 rescue the respiratory deficiency of Δ*cox12*. A) 10-fold serial dilutions of cultures of various strains exposed to a temperature of 50°C for the indicated time (0, 5, 10, 15, 20, 45 min) were deposited on YPD plates and grown for 2 days at 30°C. The data are representative of 2 independent experiments. B) 10-fold serial dilutions of control strains (WT and Δ*cox12* ancestor) and of Δ*cox12* Δ*hsp104* containing or not an integrated version of the Hsp104-A375V allele. C) 10-fold serial dilutions of the Δ*cox12* ancestor containing an empty plasmid (vec), low copy (lc) or high copy (hc) plasmids encoding Hsp104 or Hsp104-A375V.

### Deletion and overexpression of Hsp104 rescue the respiratory growth defect of Δ*cox12*

Next, we wanted to verify whether the *hsp104*-C1124T mutation is sufficient to restore LGA growth of the Δ*cox12* strain. For this, we crossed a Δ*cox12* strain to the Δ*hsp104* strain with integrated *hsp104*-C1124T and isolated double mutants from the meiotic progeny after tetrad dissection (data not shown). The Δ*cox12* Δ*hsp104* + Hsp104-A375V strain showed robust growth on respiratory medium, but surprisingly, so did the Δ*cox12* Δ*hsp104* strain (Fig. 3B). We then evaluated the effect of plasmids encoding Hsp104 or Hsp104-A375V on the respiratory phenotype of the Δ*cox12* ancestor. Low-copy plasmids allowed intermediate growth and high copy plasmids provided strong complementation, whether they carried WT Hsp104 or the A375V allele (Fig. 3C). Together, our results show that Hsp104 influences positively the respiratory capacities of the Δ*cox12* strain in multiple ways.

### Guanidium hydrochloride improves the growth of Δ*cox12* cells on respiratory medium

Besides its role in thermotolerance, Hsp104 has also been connected with the maintenance of the [*PSI+*] prion ^18,19^. [*PSI+*] is a naturally occurring amyloid prion consisting of highly ordered fibrous aggregates of the Sup35 protein ^20^. The Hsp104 disaggregase activity is required for prion replication, likely by fragmenting the fibers, thus creating new seeds for prion propagation ^18^. The overexpression of Hsp104 has also been shown to efficiently prevent the transmission of [*PSI+*] to the mitotic progeny ^21^, the widely held view being that increased levels of Hsp104 cause complete disaggregation of the Sup35 amyloid fibers ^22^. Thus, the inactivation of Hsp104 or its overexpression can both cure the [*PSI^+^*] prion from *S. cerevisiae* ^22^. Given that deletion of *hsp104* and overexpression of Hsp104 restored respiratory growth of the Δ*cox12* strain (Fig. 3B, 3C), we investigated whether this phenotype was linked to [*PSI^+^*].

Guanidine hydrochloride (Gu) is known to inactivate Hsp104 and consequently blocks the replication of prions, particularly [*PSI^+^*] ^19^. Therefore, we evaluated the effect of adding Gu to the culture medium of WT and Δ*cox12* strains prior to drop-test or directly to the solid medium used for phenotypic characterization. Gu decreased the size of the colonies on glucose medium and had a negative impact on the respiratory growth of the WT strain when added to the liquid culture (Fig. S3). Even though we noticed a mild improvement of growth of Δ*cox12* cells on YPLG+Gu compared to YPLG (Fig. S3), these experimental conditions are not adequate to evaluate a putative recovery of respiratory growth of the Δ*cox12* strain, since the toxicity of Gu might impact the phenotype. We reasoned that the toxicity should be alleviated upon withdrawal of Gu whereas the elimination of the prion would be long-lasting. Thus, we cultivated a Δ*cox12/COX12* heterodiploid strain with or without Gu, then we sporulated the cells and we dissected tetrads in the absence of Gu. Replica-plating onto YPD + G418 identified the Δ*cox12* strains and their respiratory growth was evaluated on YPLG+ G418 plates (Fig. 4A). Several large colonies were observed on the YPLG + G418 plate for cells pretreated with Gu, whereas only small colonies arose from cells not previously exposed to Gu (Fig. 4A). We selected three Δ*cox12* strains from each condition (1-6 on Fig. 4A) and assessed their growth using serial dilutions (Fig. 4B). Δ*cox12* clones originating from cells pretreated with Gu showed robust growth on respiratory media, whereas those originating from control cells grew poorly, similar to the Δ*cox12* ancestor used as control (Fig. 4B). These results show that treatment with Gu promotes a long-lasting improvement of the respiratory growth of Δ*cox12* cells, likely as a result of curing [*PSI^+^*].

**Figure 4:**
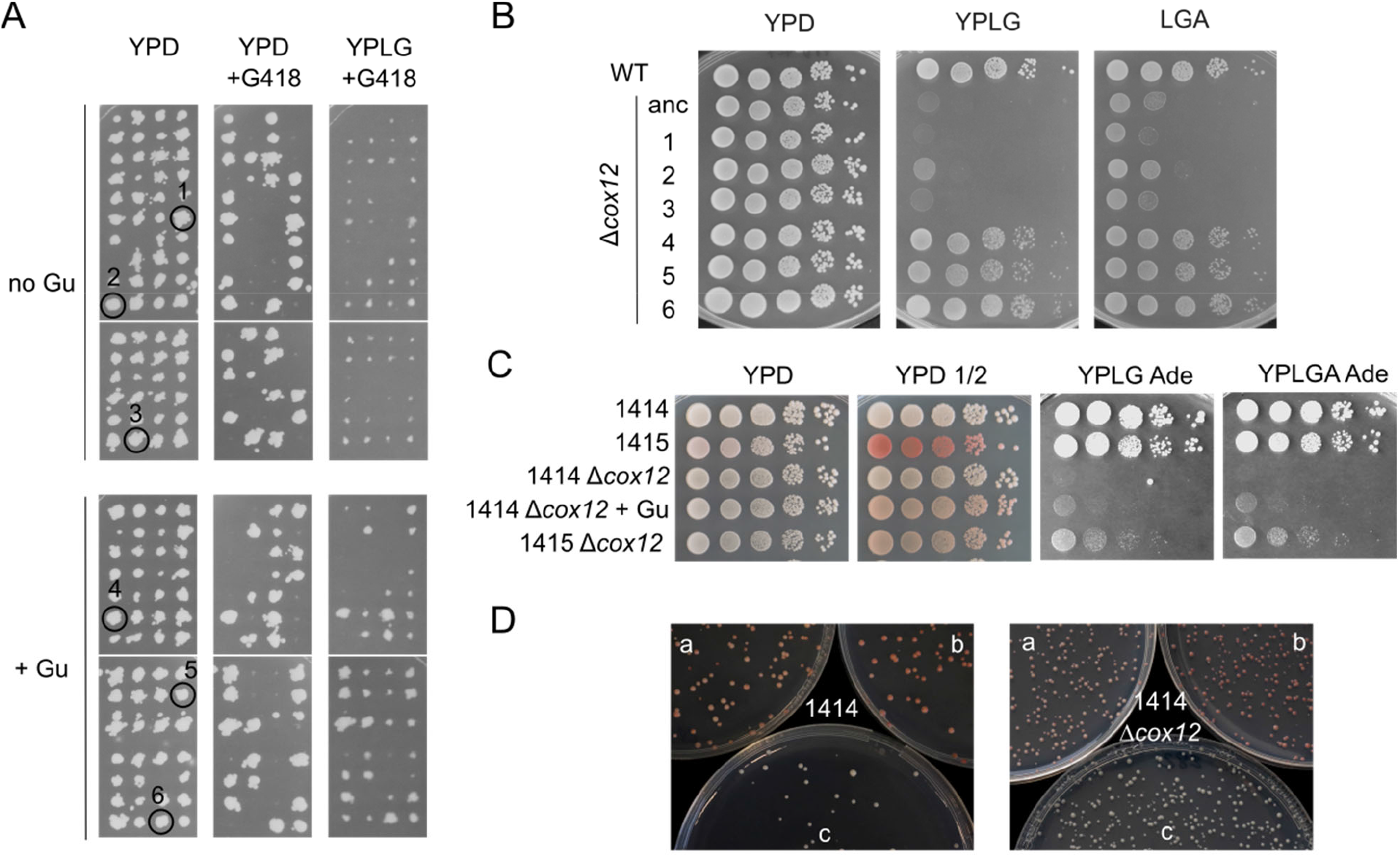
[*PSI+*] impacts negatively the respiratory growth of Δ*cox12.* A) Replica-plating of tetrads dissected from Δ*cox12*/ COX12 heterodiploid cells. Prior to sporulation, the cells were precultured twice in YPD medium containing 5 mM Gu (+Gu) or not (no Gu). B) Phenotypic analysis of the Δ*cox12* clones identified in A on YP 1% Lactate 1% Glycerol (YPLG) or synthetic medium containing 1% Lactate 0.2% Glycerol 0.2% Acetate (LGA). C) 10-fold serial dilutions of the indicated strains on YP-glucose (YPD), YP-glucose with limiting adenine (YPD ½), YP supplemented with 10X adenine and 1% Lactate 1% Glycerol (YPLG) or 1% Lactate 0.2% Glycerol 0.2% Acetate (YPLGA). +Gu indicates that the 1414Δ*cox12* strain was streaked twice onto YPD plates containing 5 mM GuHCl prior to drop-test. The plates were imaged after 3 days (YPD and YPD ½) or 8 days (YPLG and YPLGA). D) Images of colonies from 1414 or 1414Δ*cox12* strains transformed with a low copy vector encoding Hsp104 (a), Hsp104-A375V (b), or the empty vector (c) and grown for 4 days on YPD ½ plates. Results are representative of 2 independent experiments (C-D).

### The [*PSI^+^*] prion impairs mitochondrial respiration of Δ*cox12* cells

The presence of [*PSI+*] can be easily revealed by the color of cells carrying the *ade2-1* nonsense allele, which prevents the synthesis of the Ade2 protein involved in adenine biosynthesis. Indeed, *ade2-1* [*psi-*] cells devoid of the PSI prion appear red on low adenine medium due to the accumulation of a red metabolite in the adenosine pathway. On the contrary, [*PSI+*] cells appear white because the aggregation of the Sup35 protein reduces termination efficiency and causes read-through of the *ade2-1* nonsense codon ^23^, which restores adenine prototrophy and prevents the accumulation of the red metabolite ^17^. In order to directly assess the impact of [*PSI+*] on the phenotype of Δ*cox12* cells, we deleted the *COX12* gene in two isogenic backgrounds (1414 and 1415) that differ only in the presence or absence of the PSI prion ^19^. As expected, 1414 [*PSI+*] cells were white on YPD ½, whereas 1415 [*psi-*] cells appeared red (Fig. 4C). The deletion of *cox12* in strain 1415 largely attenuated the red pigmentation (Fig. 4C), consistent with the report that *pet-* cells with defective mitochondria don’t develop the red color ^24^. Importantly, 1414 Δ*cox12* cells were unable to grow on YPLG and YPLGA plates, whereas 1415 Δ*cox12* cells showed appreciable growth (Fig. 4C). Curing [*PSI+*] in 1414 Δ*cox12* cells with Gu pretreatment slightly improved the growth on respiratory media and yielded a distinctive color on YPD ½ comparable to that of 1415 Δ*cox12* cells (Fig. 4C). Overall, these data demonstrate that [*PSI+*] negatively impacts the respiratory capacity of Δ*cox12* cells.

### Hsp104-A375V cures [*PSI^+^*] efficiently

Finally, we wanted to understand the reason for the positive selection of the *hsp104*-C1124T mutation during the evolution experiment. We thus transformed low copy vectors encoding Hsp104 or Hsp104 A375V in 1414 and 1414 Δ*cox12* cells and monitored the curing of [*PSI+*] by evaluating the color of colonies on selective plates. Whereas cells containing the control vector formed white colonies as expected, most clones transformed with the Hsp104 A375V construct showed a red color (Fig. 4D). The cells encoding the WT Hsp104 protein displayed an intermediate color (Fig. 4D), suggesting that the elimination of [*PSI+*] was more efficient in cells expressing the A375V allele. Overall, we propose that the *hsp104*-C1124T mutation was selected in Δ*cox12* population B during the evolution experiment because it allowed an efficient clearing of [*PSI+*], which in turn improved CcO activity and allowed the use of the respiratory substrates available in the evolution medium.

## DISCUSSION

Classical genetic approaches, like high copy suppressor screens or the characterization of spontaneous mutants, have been instrumental to pinpoint the role of several proteins required for the proper function of the mitochondrial respiratory chain ^25–27^. However, such approaches typically yield single modifiers of a phenotype, whereas experimental evolution has the potential to reveal complex epistatic interactions involving several genes ^13,28,29^. In this study, we tested whether it is possible to compensate the defect in CcO activity caused by the absence of the peripheral subunit Cox12. Thus, we conducted an evolution experiment over several hundreds of generations to select Δ*cox12* clones with improved respiration. The two populations showed a rapid improvement of respiratory metabolism (Table S1) and genetic analysis of two representative Δ*cox12* clones showed that a single causal mutation was involved in each case. Given the high number of mutations accumulated during evolution experiments, the identification of the mutation(s) causative of the phenotype is often challenging ^13,30^. Here, we identified *hsp104*-C1124T as the causative mutation in population B via next generation sequencing of the meiotic progeny of evolved clones selected for respiratory growth on LGA medium. The reasons for our incapacity to identify the causal mutation in population A remain unclear but this mutation is interesting as it restored CcO activity and respiratory growth to levels comparable to those obtained in population B (Fig. 1–2).

We showed that cells containing the *hsp104*-C1124T allele, which encodes the Hsp104-A375V variant, lost the [*PSI+*] prion more efficiently compared to cells with wild-type Hsp104 (Fig. 4D). The importance of the AAA+ disaggregase Hsp104 for the propagation of [*PSI+*] has been recognized early on with studies showing that [*PSI+*] was cured not only by loss of function of Hsp104, but also by its overexpression ^31,32^. The same basic function of Hsp104, namely the disentangling of misfolded/aggregated proteins by extrusion through the central pore of the Hsp104 hexamer, is needed for its role in prion propagation and induced thermotolerance ^22^. As expected, many point mutations in Hsp104 that decrease thermotolerance also decrease prion propagation ^33^. Nevertheless, both functions can be dissociated since some mutations were described in Hsp104 to cause defective prion propagation despite maintaining thermotolerance ^33,34^. The A375V mutation that we identified in the NBD1 also falls into this class since it maintained thermotolerance and cured [*PSI+*] efficiently.

Several prion proteins have been identified in *S. cerevisiae*, the most studied being [*PSI+*] and [*PIN+*] ^20^. They correspond to infectious and self-replicating amyloid-like aggregates of Sup35 and Rnq1, respectively. The Sup35 protein functions normally as a translation termination factor, thus aggregation of Sup35 in [*PSI+*] cells increases the rate of ribosomal read-through for nonsense codons to ~1%, as compared to ~0.3% in [*psi-*] cells ^35^. As such, [*PSI+*] would promote evolvability and is thus thought to be beneficial in harsh environments ^36^, but its rare frequency in nature – presence in ~1% of 700 wild yeast isolates – suggests that it may be detrimental in most cases ^36^. Rates of spontaneous appearance of [*PSI+*] in laboratory conditions range from 6*10^-7^ to 1*10^-5^ per generation ^37,38^ and increase up to 4*10^-4^ during chronological aging ^37^. Even though the effect of [*PSI+*] on cellular fitness in laboratory conditions is still debated ^20,37,39^, the self-propagating properties of prions may result in a rather high frequency of [*PSI+*] in laboratory strains.

Interestingly, [*PSI+*] has been previously connected to mitochondrial function in yeast. Indeed, [*PSI+*] cells showed a highly fragmented mitochondrial network and a decreased mitochondrial abundance of the prohibitins Phb1 and Phb2, probably as a result of being trapped in cytosolic [*PSI+*] aggregates ^40^. Since prohibitins stabilize newly synthesized mitochondrial proteins ^41^, Cox2 was proposed to be indirectly destabilized by [*PSI+*] via the decrease of Phb1 and Phb2 ^40^. This could contribute to the [*PSI+*]-specific respiratory deficiency observed in the *nam9-1* strain, which carries a point mutation in the Nam9 mitoribosome subunit ^42^.

In *S. cerevisiae*, CcO is composed of 11 nuclear-encoded subunits and 3 mitochondrial-encoded subunits. Overall, our data show that the function of CcO *in vivo* is not strictly dependent on Cox12 since the Δ*cox12* ancestor displayed a ~20% residual CcO activity and the evolved Δ*cox12* clones Ba-P2 and Ac-P1 recovered ~50% activity of WT cells (Fig. 2E). These data are consistent with previous work showing that purified CcO was active despite the loss of the Cox12 subunit during the purification ^7^. Since several studies reported that CcO subunits are degraded when the enzyme is not properly assembled ^43–45^, it is tempting to speculate that the low levels of Cox1,2,13 proteins observed in Δ*cox12* cells reflect an assembly defect of CcO, which is partially corrected in the evolved clones (Fig. 2A). Interestingly, Cox12 is rapidly ubiquitinated and degraded by the proteasome when its mitochondrial import is defective ^46^. Thus, the addition of Cox12 into CcO, which corresponds to one of the last step in the assembly process ^47^, appears to be tightly controlled by cytosolic proteostatis.

The assembly of functional CcO is a complex process that requires multiple accessory factors to coordinate the synthesis, maturation and assembly of the 14 subunits ^43,47,48^. In this study, we demonstrated in two genetic backgrounds (CEN-PK2 and 1414/5) that [*PSI+*] is detrimental to the respiratory metabolism of Δ*cox12* cells and that [*PSI+*] negatively impacts the activity of CcO. Several mechanisms might underlie these phenotypes. First, the assembly of CcO might be impaired by lower levels of particular CcO subunits or assembly factors, as a result of cytosolic sequestration within Sup35 aggregates or via destabilization due to decreased abundance of stabilizing proteins like prohibitins ^40^. Even though a recent proteomic study demonstrated that the proteomes of [*psi−*] and [*PSI+*] cells are very similar ^49^, variations in subcellular levels, notably of mitochondrial proteins, are possible. Second, [*PSI+*] is known to decrease the efficiency of translation termination, which could perturb the synthesis of critical nuclear-encoded CcO subunits or assembly factors and thus impair the concerted assembly of CcO. However, the rather modest increase in read-through rate - from 0.3% in [*psi-*] cells to 1% in [*PSI+*] cells ^35^-suggests that its impact is likely limited. Whatever the mechanism at play, we clearly demonstrated in this study that [*PSI+*] impairs mitochondrial respiration in Δ*cox12* cells. Our work reveals that [*PSI+*] is an important modulator of the complex interplay between the cytosol and mitochondria. Whether [*PSI+*] affects other respiratory complexes besides CcO and whether other prions may also impact mitochondrial bioenergetics are interesting questions that remain to be addressed in future studies.

## MATERIAL AND METHODS

### Yeast culture conditions

Yeast strains were typically grown at 30°C and 180 rpm shaking in either YP-based rich medium (1% [w/v] yeast extract, 2% [w/v] peptone), or in YNB-based synthetic medium (YNB wo AS, US Biological) supplemented with ammonium sulfate (5g/L) and nutrients to cover the strains’ auxotrophies in quantities as described in ^50^. Carbon sources from 20% stocks sterilized by filtration were added at the following final concentration, unless indicated otherwise: glucose (2% [w/v]), lactate (2% [w/v]), ethanol (2% [v/v]), acetate (2% [w/v]), glycerol (2% [v/v]), LGA was 1% lactate-0.1% glycerol-0.1% acetate. When G418 was added to synthetic medium (0.2mg/mL), ammonium sulfate was omitted and monosodium glutamate (MSG, 1g/L) was used as sole nitrogen source. ½ YPD (0.5% yeast extract, 2% peptone, 2% glucose) was used to monitor the PSI status of *ade2-1* strains. Bacto Agar (Euromedex) was added at 1.6% (w/v) for solid media.

### Construction of strains and plasmids

*Saccharomyces cerevisiae* strains used in this study are listed in Table S4. Transformations were performed using the lithium acetate method ^51^ and genomic DNA was prepared according to ^52^. The Δ*cox12* strain used in the evolution experiment was constructed from the CEN.PK2-1D strain in several steps including insertion of *Saccharomyces kluyveri HIS3* gene at the *MET17* locus, restoration of *LEU2* at the *leu2-3_112* locus, deletion of *cox12* by the KanMX cassette and deletion of the DNA mismatch repair gene *msh2* by *Candida albicans URA3*. First, the *HIS3* gene from *S. kluyveri* was amplified from the pFA6a:His3MX6 plasmid using the oligonucleotides 5SkHis3Met17 and 3SkHis3Met17 (Table S5) and the PCR product was transformed in CEN.PK2-1D. Recombinant clones were selected on medium lacking histidine and were checked for methionine auxotrophy. Correct replacement of MET17 ORF by the SkHIS3 gene was verified by PCR with the 5verifMet17 and 3verifMet17 primers and one clone, called CEN-H, was selected for subsequent modification of the *leu2-3_112* locus. A PCR product corresponding to the *LEU2* gene was amplified from the pRS415 plasmid with oligonucleotides 5verifLeu2 and 3verifLeu2 and transformed into CEN-H. Recombinant clones were selected on medium lacking leucine and correction of the LEU2 locus in strain CEN-HL was verified by DNA sequencing. Alternatively, the *leu2-3_112* locus of CEN-H was replaced with the *MET17* gene by homologous recombination with a PCR fragment obtained with the primers 5Met17Leu2 and 3Met17Leu2. Recombinants were selected on medium lacking methionine and correct insertion of the *MET17* gene at the *leu2* locus was verified by PCR with 5verifleu2 and 3verifleu2 primers on the genomic DNA. One correct clone was selected and named CEN-HM. Strain CEN-HL was crossed to CEN.PK2-1C, the diploid was selected on medium lacking leucine and methionine, and after sporulation according to a published procedure ^50^, tetrads were dissected on a Nikon 50i microscope equipped with a micromanipulator. Strains CEN-HLa and CEN-Ha were isolated after selecting the appropriate markers in the meiotic progeny and verifying the loci by PCR. *COX12* was deleted in strains CEN-HLa and CEN-HM by homologous recombination of a fragment obtained by PCR amplification with the primers 5cox12 and 3cox12 on the genomic DNA of the BY4741 Δ*cox12::KanMX4* strain. Recombinants were selected on YPD medium containing G418, positive clones were then plated on YPLG to verify respiratory growth and strains Δ*cox12-HLa* and Δ*cox12-HM* were confirmed by PCR amplification with the primers 5cox12 and 3cox12. Finally, the *MSH2* gene was deleted in the Δ*cox12-HLa* strain by homologous recombination with a fragment obtained by PCR using the primers 5KOmsh2 and 3KOmsh2 on genomic DNA of Candida albicans SC5314. Transformants were selected on medium lacking uracil and correct insertion of CaURA3 at the *msh2* locus was verified by PCR with primers 5verifMsh2 and 3verifMsh2 on the genomic DNA of candidates. The resulting strain was named Δ*cox12-msHLa* and was used as the ancestral strain (Δ*cox12* anc.) to initiate the evolution experiment.

Deletion of *cox12* in the 1414 and 1415 strains (derived from the 779-6A strain ^19^) was conducted as described above with recombination of the kanMX4 cassette at the *cox12* locus. Hsp104 was deleted in the CEN.PK2-1C strain by replacing the *hsp104* locus with a URA3 cassette, amplified with primer Hsp104_fwd and Hsp104_rev from a pUG72 plasmid.

All plasmids used in this study are listed in Table S6. pRS424 plasmids expressing wild type Hsp104 or the mutant Hsp104-A375V were obtained by cloning a PCR fragment amplified from the genomic DNA of Δ*cox12* anc. or Δ*cox12* Ba-P2 with primers cHsp104_fwd and cHsp104_rev. The PCR fragment contained the *HSP104* ORF with its endogenous promoter and terminator and was cloned into pRS424 using *XhoI* and *NotI*. After verification by sequencing, inserts were subcloned into the pRS414 plasmid. Hsp104-A375V was also subcloned into a pRS305 plasmid using *XhoI* and *NotI* and subsequently integrated into Δ*hsp104* to yield Δ*hsp104*+Hsp104-A375V. Strains Δ*cox12*Δ*hsp104* and Δ*cox12*Δ*hsp104*+Hsp104-A375V were isolated from the F1 meiotic progeny of a diploid obtained by mating Δ*cox12-HM* with Δ*hsp104*+Hsp104-A375V. Diploids were selected on –leucine – histidine medium, sporulated and tetrads were dissected. Germinated spores were screened for LEU2, URA3 and kanMX4 markers, loci were verified by PCR and Δ*cox12*Δ*hsp104* (*URA3, kanMX4, leu2*) and Δ*cox12*Δ*hsp104*+Hsp104-A375V (*URA3, kanMX4, LEU2*) were obtained.

### Experimental evolution

The medium was composed of 1.67g/L YNB (without pABA and folate, MP Biomedicals) supplemented with 5g/L ammonium sulfate, 1g/L monosodium glutamate, 0,008% yeast extracts, 0,15% maltose, 1% Lactate / 0,2% Glycerol / 0,2% Acetate (LGA), tryptophan and methionine to cover the auxotrophies ^50^. The pH of the medium was adjusted to 4.4 with 5 M NaOH and the carbon sources (sterilized by filtration) were added after autoclaving. The Δ*cox12-msHLa* strain was inoculated in duplicate in 250 mL Erlenmeyer flasks containing 20 mL medium and incubated at 30°C with 180 rpm shaking. Both cultures were diluted and transferred daily into fresh medium for 28 days, (see Table S1 for dilution factors), then from day 29, the cultures were transferred only every two days. Before transfer, growth was evaluated by measuring the optical density at 600nm (Table S1) in a Tecan microplate reader (NanoQuant Infinite M200PRO). From transfer 8, maltose was omitted from the culture medium. Every 3 to 4 transfers, glycerol stocks were constituted by sampling 0.8 mL of culture, adding 0.2 mL glycerol, and freezing in liquid nitrogen before storing at −80°C.

### Genetic analysis of clones from experimental evolution

Among the three clones Aa-Ac and Ba-Bc, which we isolated from each population at the end of the evolution experiment (after growth of cells from transfer 43), Ac and Ba were selected for further analysis. To differentiate between dominant and recessive causal mutations, Ac and Ba were crossed with the Δcox12-HM strain and diploids were selected on medium lacking leucine and methionine.

To evaluate the number of causal mutation(s) in clones Ac and Ba, they were crossed to strain CEN-HM and diploids were selected on medium lacking leucine and methionine. We sporulated the diploids and performed tetrad dissection on glucose synthetic medium. After 3 days at 30°C, we replicated about 15 complete tetrads for each strain on synthetic glucose media lacking leucine or methionine, synthetic LGA medium containing or not G418, on YPD + G418. The plates were incubated for 2 days (glucose) or 4 days (LGA) at 30°C. Clones that grew on the YPD + G418 carried the Δ*cox12::kanMX4* marker and were subdivided into P clones and N clones depending on their growth phenotype on the LGA plate (P = Positive for growth, N = Negative for growth). 10-15 individual P and N clones were grouped to constitute the P and N pools used for whole genome sequencing. Ac-P1/2 and Ba-P1/2 clones were characterized phenotypically and Ac-P1, Ba-P2 were characterized biochemically (Fig. 2).

### Whole genome sequencing

Selected clones were grown in 10 mL cultures in synthetic glucose medium for 30 hours at 30°C, 180 rpm shaking. For cultures of P and N clones, 2 units OD600 were mixed together to obtain Ac-P, Ac-N, Ba-P and Ba-N pools. Cells were collected by centrifugation 3200g, 4°C, 5 min and genomic DNA was prepared from cell pellets according to ^52^. Illumina sequencing was performed by the Max Planck-Genome-centre Cologne, Germany (https://mpgc.mpipz.mpg.de/home/). Genomic DNA was sheared by Covaris Adaptive Focused Acoustics technology (COVARIS, Inc.) to average fragment size of 250bp with settings: intensity 5, duty cycle 10%, 200 cycles per burst and 180s treatment time. Library preparation was done with NEBNext Ultra DNA Library preparation kit, and then sequenced as a 2×150bp paired end reads on a HiSeq 3000 to approximately 3 million reads per sample. The sequencing data were deposited in BioProject at NCBI under the identification PRJNA768319.

### Analysis of whole genome sequencing data

Reads were analyzed with fastQC (https://www.bioinformatics.babraham.ac.uk/projects/fastqc/) and trimmed with cutadapt based on the fastQC results ^53^. Trimmed reads were aligned on the S288C R64-1-1 reference genome using bwa-mem (arXiv:1303.3997v2). The resulting BAM files were sorted and indexed with samtools ^54^. GATK HaplotypeCaller 3.8 in BP_RESOLUTION mode was used to produce GVCF files ^55^. The resulting GVCF files were merged with mergeGVCFs.pl (a component of https://github.com/ntm/grexome-TIMC-Primary) to obtain a single GVCF file. Since some samples were clonal while others were pools of 10 to 15 haploid strains, the GATK genotype calls (GT) were often irrelevant; they were therefore discarded, and the ALT alleles and allele frequencies (AF) produced by GATK were used to make our own genotype calls, using a simple yet reliable algorithm. Briefly, a minimum depth DP of 10 was imposed to make a call, and any allele supported by at least 15% of the reads was called for that sample.

Variants were then filtered, imposing a homozygous variant call in samples Ba-P2 and Ba-P pool and a homozygous reference call in all other samples. This resulted in three candidate variants (Table S3). Annotation using VEP ^56^ showed that a single variant had more than “MODIFIER” impact, and directly affected a coding sequence: the XII:89746 C→T substitution, which results in a A375V missense mutation in HSP104. Similar analyses with the Ac pool and Ac-P1 clone did not yield any promising candidates. All scripts developed for these analyses are available (https://github.com/ntm/cox12_evolutionExperiment).

### Growth assay by serial dilution

Overnight cultures were adjusted in sterile water to OD_600_ = 1 and ten-fold serial dilutions were performed in sterile water in 96 well plates. 5 μL of each dilution was spotted on various plates using multi-channel pipette and plates were typically imaged after incubation at 30 °C for three days (fermentable carbon source) or 5 to 10 days (respiratory carbon sources).

### Induced thermotolerance assay

Cells from overnight cultures were used to inoculate 4 mL YPD cultures at OD_600nm_ = 0.2. The cultures in glass tubes were incubated at 30°C to mid log phase (OD_600nm_~1) at which point they were switched to 37°C for one hour with 180 rpm shaking. Then, cultures were transferred at 50°C in a water bath with occasional shaking and 200 μL aliquots were taken at different time intervals (3, 7, 10, 15, 20, 30 and 45 min), diluted in series (5 dilutions of 10 fold each) in 96 well plates. 5 μL of each dilution were spotted on a YPD agar plate and the plate was incubated for 2 days at 30°C.

### Immunoblotting

To determine steady state protein levels, strains were grown either on YPD or YPGal to mid-log phase and cells equivalent to an OD_600_ of 3 were harvested. Pellets were resuspended in 200 μL of 0.1 M NaOH and incubated at room temperature (RT) for 5 min with 1400 rpm shaking. Samples were centrifuged (1500 rcf, 5 min, RT) and pellets were resuspended in 50 μL of reducing Laemmli buffer (50 mM Tris-HCl, 2% SDS, 10% glycerol, 0.1% bromophenol blue, 100 mM DTT; adjusted to pH 6.8). Subsequently, samples were incubated for 5 min at RT, 1400 rpm shaking and heated for 10 min to 65°C prior to SDS-PAGE. 10 μL of the sample were applied for standard SDS-PAGE and immunoblotting followed standard protocols. Blots were decorated with antibodies against Tom70, Cor1, Cor2, Cox1, Cox2, Cox4, Cox12, Cox13 and Rcf2 as described previously ^57^.

### Mitochondrial isolation

Mitochondria were isolated according to ^58^. In brief, yeast cells were grown in YPGly media to mid-log phase and harvested by centrifugation (3000 rcf, 5 min, RT). Cells were washed once in distilled water and resuspended in 2 mL/g cell wet weight MP1 buffer (0.1 M Tris, 10 mM dithiothreitol, pH 9.4). After incubation for 10 min at 30°C, 170 rpm shaking, cells were washed once in 1.2 M sorbitol and subsequently resuspended in 6.7 mL/ cell wet weight MP2 buffer (20 mM potassium phosphate, 0.6 M sorbitol, pH 7.4, containing 3 mg/g of cell wet weight zymolyase 20T). Spheroplasts were created via incubation for 1 h at 30°C and harvested by centrifugation (3000 rcf, 5 min, 4°C). After careful resuspension in 13.4 mL/g of cell wet weight in ice-cold homogenization buffer (10 mM Tris, 0.6 M sorbitol, 1 mM ethylenediaminetetraacetic acid, 1 mM phenylmethylsulfonyl fluoride, pH 7.4), lysates were created by mechanical disruption via ten strokes with a Teflon plunger. Homogenates were centrifuged for 5 min at 3000 rcf, 4°C and the resulting supernatants were subsequently centrifuged at 17000 rcf for 12 min, 4°C. Pelleted mitochondria were resuspended in isotonic buffer (20 mM HEPES, 0.6 M sorbitol, pH 7.4) to a concentration of 10 mg/mL and stored at −80°C.

### UV-VIS spectroscopy and CcO activity

A Cary4000 UV-VIS spectrophotometer (Agilent Technologies) was used to record optical spectra (400-650 nm) from isolated mitochondria. 200 μg of mitochondria were lysed in 20 μL of lysis buffer (50 mM KPi pH 7.4, 150 mM KCl, 1× Complete Protease Inhibitor cocktail (Roche), 1 mM PMSF, 1% n-Dodecyl β-D-maltoside) for 20 min at 4°C and subsequently mixed with 130 μL of dilution buffer (50 mM KPi pH 7.4, 150 mM KCl) in a microcuvette and applied for measurement. To obtain reduced spectra, small amounts of sodium dithionite were admixed and measurements were repeated. Heme concentrations were determined from the differential spectrum (reduced-minus-oxidized) from 2 replicates by applying calculations as described recently ^57^. Concentrations of CcO were determined by applying an extinction coefficient of ∊ = 26 mM-1 cm-1 and volumes of isolated mitochondria corresponding to 10 nmol CcO were used for CcO activity measurements. Thereby, mitochondria were lysed in lysis buffer (50 mM Tris pH 7.4, 100 mM KCl, 1 mM EDTA, 1× Complete Protease Inhibitor cocktail (Roche), 1mM PMSF, 2% Digitonin) for 10 min, 4°C and clarified by centrifugation (25000 rcf, 10 min, 4°C). Lysates were diluted in measurement buffer (50 mM Tris pH 7.4, 100 mM KCl, 1 mM EDTA) and transferred into a Clark-type oxygen electrode. The measurement was started by adding 20 mM Na-Ascorbate, 50 μM Yeast Cyt. *c*, 40 μM N,N,N′,N′-tetramethyl-p-phenylenediamine (TMPD) and recorded for 1-2 min. The slope of each analysis as a measure for oxygen consumption (μmol O2/ml/min) was evaluated from 3 replicates and normalized to the average of the slope from wild type mitochondrial lysates to describe CcO activity as fold change.

### Statistical analysis

Outliers were defined as data points outside 1.5-fold interquartile range and of note, no outliers were detected with this method. Normal distribution of the data was confirmed by a Shapiro-Wilk’s test (R studio, shapiro_test) and homogeneity of variances by a Leven’s test (R studio, leveneTest). The means were compared by a One-Way ANOVA followed by a Tukey post hoc test (R studio, anova_test, tukey_hsd). Significances are indicated with asterisk (***P < 0.001, **P < 0.01, *P < 0.05, n.s. P > 0.05) and graphs were created via R studio using the ggplot2 library.

## ACKNOWLEDGMENTS

This work was supported by CNRS and Université Grenoble-Alpes (to FP) and by the Swedish Research Council (2018-03694 to MO), the Knut and Alice Wallenberg foundation (2013.0006 to MO). Andreas Aufschnaiter was supported by an Erwin Schrödinger Fellowship from the Austrian Science Fund FWF (J4398-B). The group of James Bruce Stewart was supported by the Max Planck Society. We thank the Max Planck-Genome-centre Cologne for performing the Illumina sequencing in this study, and support from the Bioinformatics Core facility at the Max Planck for Biology of Ageing. We thank Muriel Cornet for providing genomic DNA of *Candida albicans* SC5314, and Dan Masison (NIH, Bethesda) for providing the 1414 and 1415 strains and for critical reading of the manuscript.

## REFERENCES

(1) Pfanner, N.; Warscheid, B.; Wiedemann, N. Mitochondrial Proteins: From Biogenesis to Functional Networks. Nat Rev Mol Cell Biol 2019, 20 (5), 267–284. https://doi.org/10.1038/s41580-018-0092-0.

(2) Couvillion, M. T.; Soto, I. C.; Shipkovenska, G.; Churchman, L. S. Synchronized Mitochondrial and Cytosolic Translation Programs. Nature 2016, 533 (7604), 499–503. https://doi.org/10.1038/nature18015.

(3) Ott, M.; Amunts, A.; Brown, A. Organization and Regulation of Mitochondrial Protein Synthesis. Annu Rev Biochem 2016, 85, 77–101. https://doi.org/10.1146/annurev-biochem-060815-014334.

(4) Carr, H. S.; Winge, D. R. Assembly of Cytochrome c Oxidase within the Mitochondrion. Acc Chem Res 2003, 36 (5), 309–316. https://doi.org/10.1021/ar0200807.

(5) van der Sluis, E. O.; Bauerschmitt, H.; Becker, T.; Mielke, T.; Frauenfeld, J.; Berninghausen, O.; Neupert, W.; Herrmann, J. M.; Beckmann, R. Parallel Structural Evolution of Mitochondrial Ribosomes and OXPHOS Complexes. Genome Biol Evol 2015, 7 (5), 1235–1251. https://doi.org/10.1093/gbe/evv061.

(6) Rathore, S.; Berndtsson, J.; Marin-Buera, L.; Conrad, J.; Carroni, M.; Brzezinski, P.; Ott, M. Cryo-EM Structure of the Yeast Respiratory Supercomplex. Nat Struct Mol Biol 2019, 26 (1), 50–57. https://doi.org/10.1038/s41594-018-0169-7.

(7) LaMarche, A. E.; Abate, M. I.; Chan, S. H.; Trumpower, B. L. Isolation and Characterization of COX12, the Nuclear Gene for a Previously Unrecognized Subunit of Saccharomyces Cerevisiae Cytochrome c Oxidase. The Journal of biological chemistry 1992, 267 (31), 22473–22480.

(8) Brischigliaro, M.; Zeviani, M. Cytochrome c Oxidase Deficiency. Biochim. Biophys. Acta-Bioenerg. 2021, 1862 (1), 148335. https://doi.org/10.1016/j.bbabio.2020.148335.

(9) Ghosh, A.; Pratt, A. T.; Soma, S.; Theriault, S. G.; Griffin, A. T.; Trivedi, P. P.; Gohil, V. M. Mitochondrial Disease Genes COA6, COX6B and SCO2 Have Overlapping Roles in COX2 Biogenesis. Hum. Mol. Genet. 2016, 25 (4), 660–671. https://doi.org/10.1093/hmg/ddv503.

(10) Gammie, A. E.; Erdeniz, N.; Beaver, J.; Devlin, B.; Nanji, A.; Rose, M. D. Functional Characterization of Pathogenic Human MSH2 Missense Mutations in Saccharomyces Cerevisiae. Genetics 2007, 177 (2), 707–721. https://doi.org/10.1534/genetics.107.071084.

(11) Stone, J. E.; Petes, T. D. Analysis of the Proteins Involved in the in Vivo Repair of Base-Base Mismatches and Four-Base Loops Formed during Meiotic Recombination in the Yeast Saccharomyces Cerevisiae. Genetics 2006, 173 (3), 1223–1239. https://doi.org/10.1534/genetics.106.055616.

(12) Codon, A. C.; Gasentramirez, J. M.; Benitez, T. FACTORS WHICH AFFECT THE FREQUENCY OF SPORULATION AND TETRAD FORMATION IN SACCHAROMYCES-CEREVISIAE BAKERS YEASTS. Appl. Environ. Microbiol. 1995, 61 (2), 630–638.

(13) Koschwanez, J. H.; Foster, K. R.; Murray, A. W. Improved Use of a Public Good Selects for the Evolution of Undifferentiated Multicellularity. eLife 2013, 2, 27. https://doi.org/10.7554/eLife.00367.

(14) Glover, J. R.; Lindquist, S. Hsp104, Hsp70, and Hsp40: A Novel Chaperone System That Rescues Previously Aggregated Proteins. Cell 1998, 94 (1), 73–82. https://doi.org/10.1016/s0092-8674(00)81223-4.

(15) Hattendorf, D. A.; Lindquist, S. L. Cooperative Kinetics of Both Hsp104 ATPase Domains and Interdomain Communication Revealed by AAA Sensor-1 Mutants. EMBO J 2002, 21 (1–2), 12–21. https://doi.org/10.1093/emboj/21.1.12.

(16) Sanchez, Y.; Lindquist, S. L. HSP104 REQUIRED FOR INDUCED THERMOTOLERANCE. Science 1990, 248 (4959), 1112–1115. https://doi.org/10.1126/science.2188365.

(17) Hung, G. C.; Masison, D. C. N-Terminal Domain of Yeast Hsp104 Chaperone Is Dispensable for Thermotolerance and Prion Propagation but Necessary for Curing Prions by Hsp104 Overexpression. Genetics 2006, 173 (2), 611–620. https://doi.org/10.1534/genetics.106.056820.

(18) Kryndushkin, D. S.; Alexandrov, I. M.; Ter-Avanesyan, M. D.; Kushnirov, V. V. Yeast [PSI+] Prion Aggregates Are Formed by Small Sup35 Polymers Fragmented by Hsp104. J. Biol. Chem. 2003, 278 (49), 49636–49643. https://doi.org/10.1074/jbc.M307996200.

(19) Jung, G.; Masison, D. C. Guanidine Hydrochloride Inhibits Hsp104 Activity In Vivo: A Possible Explanation for Its Effect in Curing Yeast Prions. Current Microbiology 2001, 43 (1), 7–10. https://doi.org/10.1007/s002840010251.

(20) Kelly, A. C.; Wickner, R. B. Saccharomyces Cerevisiae. Prion 2013, 7 (3), 215–220. https://doi.org/10.4161/pri.24845.

(21) Derkatch, I. L.; Chernoff, Y. O.; Kushnirov, V. V.; Inge-Vechtomov, S. G.; Liebman, S. W. Genesis and Variability of [PSI] Prion Factors in Saccharomyces Cerevisiae. Genetics 1996, 144 (4), 1375–1386.

(22) Greene, L. E.; Saba, F.; Silberman, R. E.; Zhao, X. Mechanisms for Curing Yeast Prions. Int. J. Mol. Sci. 2020, 21 (18), 6536. https://doi.org/10.3390/ijms21186536.

(23) Yeast Genetic Structures and Functions. In Yeast; John Wiley & Sons, Ltd, 2012; pp 73–125. https://doi.org/10.1002/9783527659180.ch5.

(24) Bharathi, V.; Girdhar, A.; Prasad, A.; Verma, M.; Taneja, V.; Patel, B. K. Use of *Ade1* and *Ade2* Mutations for Development of a Versatile Red/White Colour Assay of Amyloid-Induced Oxidative Stress in *Saccharomyces Cerevisiae*: Use of Ade1 and Ade2 Mutations for Development of Colour Assay. Yeast 2016, 33 (12), 607–620. https://doi.org/10.1002/yea.3209.

(25) Lasserre, J. P.; Dautant, A.; Aiyar, R. S.; Kucharczyk, R.; Glatigny, A.; Tribouillard-Tanvier, D.; Rytka, J.; Blondel, M.; Skoczen, N.; Reynier, P.; Pitayu, L.; Rotig, A.; Delahodde, A.; Steinmetz, L. M.; Dujardin, G.; Procaccio, V.; di Rago, J. P. Yeast as a System for Modeling Mitochondrial Disease Mechanisms and Discovering Therapies. Disease models & mechanisms 2015, 8 (6), 509–526. https://doi.org/10.1242/dmm.020438.

(26) Rutter, J.; Hughes, A. L. Power(2): The Power of Yeast Genetics Applied to the Powerhouse of the Cell. Trends Endocrinol. Metab. 2015, 26 (2), 59–68. https://doi.org/10.1016/j.tem.2014.12.002.

(27) Tzagoloff, A.; Dieckmann, C. L. Pet Genes of Saccharomyces Cerevisiae. Microbiol. Rev. 1990, 54 (3), 211–225.

(28) Kawecki, T. J.; Lenski, R. E.; Ebert, D.; Hollis, B.; Olivieri, I.; Whitlock, M. C. Experimental Evolution. Trends in ecology & evolution 2012, 27 (10), 547–560. https://doi.org/10.1016/j.tree.2012.06.001.

(29) Plucain, J.; Hindre, T.; Le Gac, M.; Tenaillon, O.; Cruveiller, S.; Medigue, C.; Leiby, N.; Harcombe, W. R.; Marx, C. J.; Lenski, R. E.; Schneider, D. Epistasis and Allele Specificity in the Emergence of a Stable Polymorphism in Escherichia Coli. Science 2014, 343 (6177), 1366–1369. https://doi.org/10.1126/science.1248688.

(30) Fisher, K. J.; Lang, G. I. Experimental Evolution in Fungi: An Untapped Resource. Fungal Genet. Biol. 2016, 94, 88–94. https://doi.org/10.1016/j.fgb.2016.06.007.

(31) Chernoff, Y.; Lindquist, S.; Ono, B.; Ingevechtomov, S.; Liebman, S. Role of the Chaperone Protein Hsp104 in Propagation of the Yeast Prion-Like Factor [Psi(+)]. Science 1995, 268 (5212), 880–884. https://doi.org/10.1126/science.7754373.

(32) Wegrzyn, R. D.; Bapat, K.; Newnam, G. P.; Zink, A. D.; Chernoff, Y. O. Mechanism of Prion Loss after Hsp104 Inactivation in Yeast. Mol. Cell. Biol. 2001, 21 (14), 4656–4669. https://doi.org/10.1128/MCB.21.14.4656-4669.2001.

(33) Kurahashi, H.; Nakamura, Y. Channel Mutations in Hsp104 Hexamer Distinctively Affect Thermotolerance and Prion-Specific Propagation. Mol. Microbiol. 2007, 63 (6), 1669–1683. https://doi.org/10.1111/j.1365-2958.2007.05629.x.

(34) Jung, G.; Jones, G.; Masison, D. C. Amino Acid Residue 184 of Yeast Hsp104 Chaperone Is Critical for Prion-Curing by Guanidine, Prion Propagation, and Thermotolerance. Proc Natl Acad Sci U S A 2002, 99 (15), 9936–9941. https://doi.org/10.1073/pnas.152333299.

(35) Firoozan, M.; Grant, C.; Duarte, J.; Tuite, M. Quantitation of Readthrough of Termination Codons in Yeast Using a Novel Gene Fusion Assay. Yeast 1991, 7 (2), 173–183. https://doi.org/10.1002/yea.320070211.

(36) Halfmann, R.; Jarosz, D. F.; Jones, S. K.; Chang, A.; Lancaster, A. K.; Lindquist, S. Prions Are a Common Mechanism for Phenotypic Inheritance in Wild Yeasts. Nature 2012, 482 (7385), 363-U1507. https://doi.org/10.1038/nature10875.

(37) Speldewinde, S. H.; Grant, C. M. The Frequency of Yeast [PSI+] Prion Formation Is Increased during Chronological Ageing. Microb. Cell 2017, 4 (4), 127–132. https://doi.org/10.15698/mic2017.04.568.

(38) Lancaster, A. K.; Bardill, J. P.; True, H. L.; Masel, J. The Spontaneous Appearance Rate of the Yeast Prion [PSI plus] and Its Implications for the Evolution of the Evolvability Properties of the [PSI plus] System. Genetics 2010, 184 (2), 393–400. https://doi.org/10.1534/genetics.109.110213.

(39) Kelly, A. C.; Shewmaker, F. P.; Kryndushkin, D.; Wickner, R. B. Sex, Prions, and Plasmids in Yeast. Proc. Natl. Acad. Sci. U. S. A. 2012, 109 (40), E2683–E2690. https://doi.org/10.1073/pnas.1213449109.

(40) Sikora, J.; Towpik, J.; Graczyk, D.; Kistowski, M.; Rubel, T.; Poznanski, J.; Langridge, J.; Hughes, C.; Dadlez, M.; Boguta, M. Yeast Prion [PSI+] Lowers the Levels of Mitochondrial Prohibitins. Biochim. Biophys. Acta-Mol. Cell Res. 2009, 1793 (11), 1703–1709. https://doi.org/10.1016/j.bbamcr.2009.08.003.

(41) Nijtmans, L. G. J.; Sanz, M. A.; Grivell, L. A.; Coates, P. J. The Mitochondrial PHB Complex: Roles in Mitochondrial Respiratory Complex Assembly, Ageing and Degenerative Disease. Cell. Mol. Life Sci. 2002, 59 (1), 143–155. https://doi.org/10.1007/s00018-002-8411-0.

(42) Chacinska, A.; Boguta, M.; Krzewska, J.; Rospert, S. Prion-Dependent Switching between Respiratory Competence and Deficiency in the Yeast Nam9-1 Mutant. Molecular and cellular biology 2000, 20 (19), 7220–7229.

(43) Soto, I. C.; Fontanesi, F.; Liu, J. J.; Barrientos, A. Biogenesis and Assembly of Eukaryotic Cytochrome c Oxidase Catalytic Core. Biochim. Biophys. Acta-Bioenerg. 2012, 1817 (6), 883–897. https://doi.org/10.1016/j.bbabio.2011.09.005.

(44) Barrientos, A.; Gouget, K.; Horn, D.; Soto, I. C.; Fontanesi, F. Suppression Mechanisms of COX Assembly Defects in Yeast and Human: Insights into the COX Assembly Process. Biochim. Biophys. Acta-Mol. Cell Res. 2009, 1793 (1), 97–107. https://doi.org/10.1016/j.bbamcr.2008.05.003.

(45) Zee, J. M.; Glerum, D. M. Defects in Cytochrome Oxidase Assembly in Humans: Lessons from Yeast. Biochem Cell Biol 2006, 84 (6), 859–869. https://doi.org/10.1139/o06-201.

(46) Kowalski, L.; Bragoszewski, P.; Khmelinskii, A.; Glow, E.; Knop, M.; Chacinska, A. Determinants of the Cytosolic Turnover of Mitochondrial Intermembrane Space Proteins. BMC Biol 2018, 16 (1), 66. https://doi.org/10.1186/s12915-018-0536-1.

(47) Timón-Gómez, A.; Nývltová, E.; Abriata, L. A.; Vila, A. J.; Hosler, J.; Barrientos, A. Mitochondrial Cytochrome c Oxidase Biogenesis: Recent Developments. Seminars in Cell & Developmental Biology 2018, 76, 163–178. https://doi.org/10.1016/j.semcdb.2017.08.055.

(48) Watson, S. A.; McStay, G. P. Functions of Cytochrome c Oxidase Assembly Factors. Int J Mol Sci 2020, 21 (19), E7254. https://doi.org/10.3390/ijms21197254.

(49) Chan, P. H. W.; Lee, L.; Kim, E.; Hui, T.; Stoynov, N.; Nassar, R.; Moksa, M.; Cameron, D. M.; Hirst, M.; Gsponer, J.; Mayor, T. The [PSI+] Yeast Prion Does Not Wildly Affect Proteome Composition Whereas Selective Pressure Exerted on [PSI+] Cells Can Promote Aneuploidy. Sci Rep 2017, 7, 8442. https://doi.org/10.1038/s41598-017-07999-8.

(50) Sherman, F. Getting Started with Yeast. Methods in Enzymology 2002, 350, 3–41.

(51) Burke, D.; Dawson, D.; Stearns, T. In Methods in Yeast Genetics; Cold Spring Harbor Laboratory Press, Plainview, NY, 2000.

(52) Dymond, J. S. Chapter Twelve - Preparation of Genomic DNA from Saccharomyces Cerevisiae. In Methods in Enzymology; Lorsch, J., Ed.; Laboratory Methods in Enzymology: DNA; Academic Press, 2013; Vol. 529, pp 153–160. https://doi.org/10.1016/B978-0-12-418687-3.00012-4.

(53) Martin, M. Cutadapt Removes Adapter Sequences from High-Throughput Sequencing Reads. EMBnet.journal 2011, 17 (1), 3. https://doi.org/10.14806/ej.17.1.200.

(54) Li, H.; Handsaker, B.; Wysoker, A.; Fennell, T.; Ruan, J.; Homer, N.; Marth, G.; Abecasis, G.; Durbin, R.; Genome Project Data Processing, S. The Sequence Alignment/Map Format and SAMtools. Bioinformatics 2009, 25 (16), 2078–2079. https://doi.org/10.1093/bioinformatics/btp352.

(55) McKenna, A.; Hanna, M.; Banks, E.; Sivachenko, A.; Cibulskis, K.; Kernytsky, A.; Garimella, K.; Altshuler, D.; Gabriel, S.; Daly, M.; DePristo, M. A. The Genome Analysis Toolkit: A MapReduce Framework for Analyzing next-Generation DNA Sequencing Data. Genome research 2010, 20 (9), 1297–1303. https://doi.org/10.1101/gr.107524.110.

(56) McLaren, W.; Gil, L.; Hunt, S. E.; Riat, H. S.; Ritchie, G. R. S.; Thormann, A.; Flicek, P.; Cunningham, F. The Ensembl Variant Effect Predictor. Genome Biol. 2016, 17, 14. https://doi.org/10.1186/s13059-016-0974-4.

(57) Berndtsson, J.; Aufschnaiter, A.; Rathore, S.; Marin-Buera, L.; Dawitz, H.; Diessl, J.; Kohler, V.; Barrientos, A.; Büttner, S.; Fontanesi, F.; Ott, M. Respiratory Supercomplexes Enhance Electron Transport by Decreasing Cytochrome c Diffusion Distance. EMBO Rep 2020, 21 (12), e51015. https://doi.org/10.15252/embr.202051015.

(58) Meisinger, C.; Pfanner, N.; Truscott, K. N. Isolation of Yeast Mitochondria. Methods in molecular biology 2006, 313, 33–39. https://doi.org/10.1385/1-59259-958-3:033.

