## Supplementary figures for "The [PSI+] prion and HSP104 modulate cytochrome *c* oxidase deficiency caused by deletion of COX12"

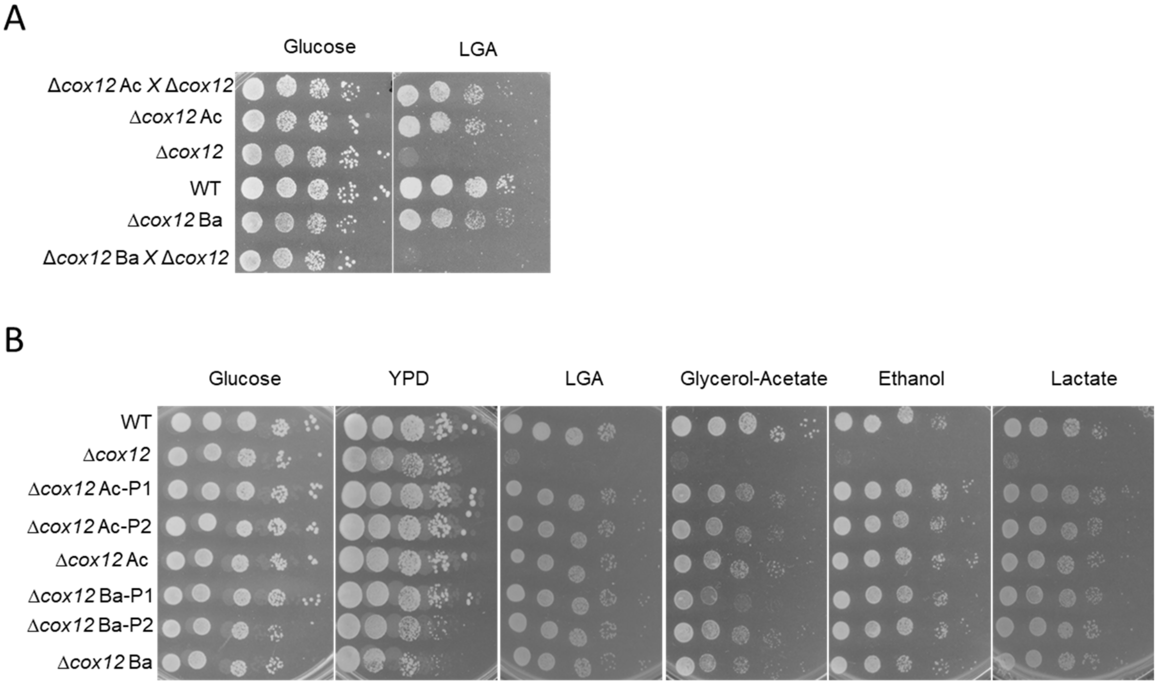


Figure S1: Genetic characterization of causal mutations in evolved Δ*cox12* cells. A) 10-fold serial dilutions of control strains (WT, Δ*cox12* ancestor, Δ*cox12* clones Ba and Ac) and of diploid strains resulting from the cross of clones Ac and Ba with the Δ*cox12-HM* strain. B) 10-fold serial dilutions of control strains (WT, Δ*cox12* ancestor, Δ*cox12* clones Ba and Ac) and of clones isolated from the meiotic progeny of the diploid strains shown in panel S1A. The plates contained either rich YPD medium or synthetic medium supplemented with the indicated carbon sources. The plates were imaged after 2 days (YPD and Glucose) or 4 days at 30°C and the data are representative of 2 independent experiments (A-B).

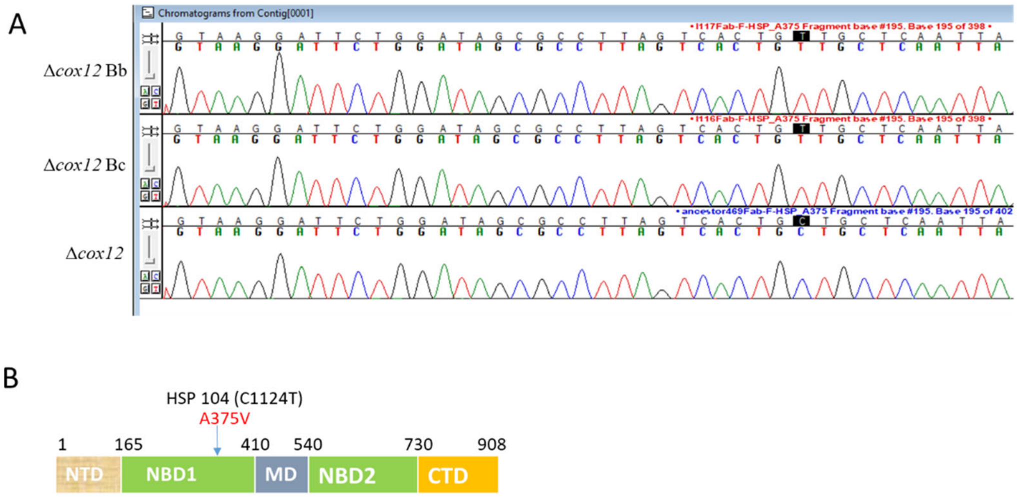


C

```
C.albicans      IQPKYEIHHGVRILDSALVTAAQLASRYLTYRALPDSAVDLVDESAAAVAVARDSKPEEL    412
S.cerevisiae    LQPKYEIHHGVRILDSALVTAAQLAKRYLPYRRLPDSALDLVDISCAGVAVARDSKPEEL    414
A.thaliana      LKEKYEGHHGVRIQDRALINAAQLSARYITGRHLPDKAIDLVDEACANVRVQLDSQPEEI    413
E.coli          LKERYELHHHVQITDPAIVAAATLSHRYIADRQLPDKAIDLIDEAASSIRMQIDSKPEEL    412
                :: :** ** *:* * *:: ** *: **:  * ***.*:**:* :.: : :  **:***:
```

Figure S2: A) Sanger sequencing chromatograms showing the presence of the C1124T mutation in PCR products amplified from Δ*cox12* clones Bb and Bc and the presence of the wild-type allele in the Δ*cox12* ancestor. B) Schematic representation of the domain composition of Hsp104 from *S. cerevisiae*. NTD (N terminal domain), NBD1 (Nucleotide binding domain 1), MD (Middle domain), NBD2 (Nucleotide binding domain 2), CTD (C-terminal domain). C) Sequence alignment of HSP104 orthologs showing the conservation of A375 (highlighted in red). The alignment was performed with Clustal Omega and the sequences are: *Arabidopsis thaliana* Hsp101 (NP_565083.1) [1], *Candida albicans* Hsp104 (AAK60626.1) , *Escherichia coli* ClpB (VWQ04789.1) [2], *Saccharomyces cerevisiae* Hsp104 (CAA97475.1).

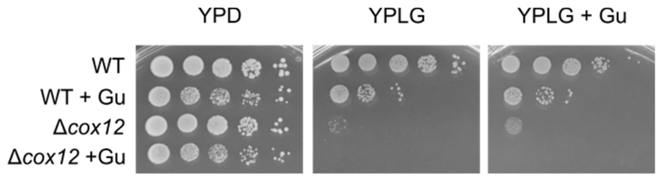


Figure S3: 10-fold serial dilutions of WT and Δ*cox12* ancestor on agar plates with YP-glucose (YPD), YP-Lactate-Glycerol (YPLG) or YPLG containing 5 mM Guanidine Hydrochloride (YPLG+ Gu). Prior to serial dilution, the strains were cultivated in liquid YPD containing or not 5 mM GuHCl.

### REFERENCES

1 Schirmer EC, Lindquist S & Vierling E (1994) An Arabidopsis heat shock protein complements a thermotolerance defect in yeast. *Plant Cell* **6**, 1899–1909.

2 Parsell DA, Sanchez Y, Stitzel JD & Lindquist S (1991) Hsp104 is a highly conserved protein with two essential nucleotide-binding sites. *Nature* **353**, 270–273.
